# Recruitment of Peroxin14 to lipid droplets affects triglyceride storage in Drosophila

**DOI:** 10.1101/2021.07.02.450950

**Authors:** Matthew N Anderson-Baron, Kazuki Ueda, Julie Haskins, Sarah C Hughes, Andrew J Simmonds

**Affiliations:** Department of Cell Biology, Faculty of Medicine and Dentistry, University of Alberta. Edmonton, AB, Canada T6G 2H7; Future Fields Ltd. Edmonton, AB, Canada; Department of Medical Genetics, Faculty of Medicine and Dentistry, University of Alberta. Edmonton, AB, Canada T6G 2H7

## Abstract

The activity of multiple organelles must be coordinated to ensure cellular lipid homeostasis. This includes the peroxisomes which metabolise certain lipids and lipid droplets which act as neutral lipid storage centres. Direct organellar contact between peroxisomes and lipid droplets has been observed, and interaction between proteins associated with the membranes of these organelles has been shown, but the functional role of these interactions is not clear. In Drosophila cells, we identified a novel localization of a subset of three transmembrane Peroxin proteins (Peroxin3, Peroxin13, and Peroxin14), normally required for peroxisome biogenesis, to newly formed lipid droplets. This event was not linked to significant changes in peroxisome size or number, nor was recruitment of other Peroxin proteins or mature peroxisomes observed. The presence of these Peroxin proteins at lipid droplets influences their function as changes in the relative levels of Peroxin14 associated with the lipid droplet surface directly affected the presence of regulatory perilipin and lipases with corresponding effects on triglyceride storage.

**Summary statement:** Interactions between peroxisomes and lipid droplets is thought to help coordinate management of cellular lipids. Peroxin proteins are required for peroxisome biogenesis. A spectrum of effects on triacylglyceride storage was seen when each of the 12 conserved Peroxins are knocked down in the *Drosophila* fat body with Peroxin14 knockdown having the largest effect. When *Drosophila* S2 cells were cultured in excess oleic acid, Peroxin3, Peroxin13, and Peroxin14, but not other Peroxins were localized to lipid droplets independently of other peroxisome markers. The presence of Peroxin14 at the lipid droplet surface altered recruitment of perilipin and lipase proteins.

## Introduction

Peroxisomes and lipid droplets (LDs) both play crucial roles in regulating cellular lipids (Lodhi and Semenkovich, 2014; Thiam and Dugail, 2019). Peroxisomes are responsible for catabolism of branched chain and very-long chain fatty acids (VLCFAs), biosynthesis of ether lipids, as well as regulating reactive oxygen (Mast et al., 2020). Structurally, peroxisomes consist of a bilayer membrane containing peroxisome membrane proteins (PMPs), surrounding a core of enzymes. LDs have a single phospholipid layer surrounding a core primarily composed of neutral lipids like triglycerides (TG) and cholesterol esters (Olzmann and Carvalho, 2019). LD activity is regulated by association of proteins like lipases and associated regulatory proteins regulating transition from TG storage to release of fatty acids (Walther et al., 2017). LDs are very large and stable in adipocytes, but smaller and transient LDs are seen in other cell types (Fujimoto and Parton, 2011). LDs form between the ER membrane leaflets and can remain connected to the ER membrane (Walther et al., 2017). Notably, yeast peroxisomes and LDs can arise from adjacent domains of the ER (Joshi et al., 2018) suggesting coordinated biogenesis (Joshi and Cohen, 2019).

In animal cells, peroxisomes are needed for β–oxidation of VLCFAs, although they can catabolize smaller-chain fatty acids when mitochondria are compromised (Violante et al., 2019). Peroxisome number, size, and composition vary based on cellular demand (Honsho et al., 2016). Peroxisomes proliferate via fission of existing peroxisomes but can also be assembled *de novo*. Either of these processes requires a source of new membrane, supplied by pre-peroxisomal vesicles (PPVs) as well as a conserved group of PMPs in the Peroxin (Pex) family. These promote recruitment of enzymes from the cytosol into the peroxisome matrix (Kim, 2017). In animal cells, PPV budding from the ER requires Pex3 and Pex16 (Fakieh et al., 2013; Geuze et al., 2003; van der Zand et al., 2010; van der Zand and Tabak, 2013). Mitochondrial-derived PPVs can also contribute to peroxisomes in a Pex3 dependent manner (Kim, 2017; Rucktaschel et al., 2010; Sugiura et al., 2017). Pex19 acts to recruit PMPs from the cytosol for insertion into the peroxisome (or PPV) membrane. Critical PMPs include Pex13 and Pex14, which form a transmembrane pore (docking complex) through which enzymes are recruited from the cytoplasm (Kim and Hettema, 2015).

LDs form within the cell when fatty acids are combined by an enzyme cascade into neutral lipids, most commonly TG, which are inserted between the ER membrane leaflets. In animal cells, these can bud from the ER and continue to grow (Walther et al., 2017). TG lipolysis releases fatty acids or other lipid species via the activity of specific lipases acting at the LD surface. Adipose triglyceride lipase (ATGL) cleaves the first fatty acyl chain from TG leaving diacylglycerol (DG) (Zimmermann et al., 2004). Hormone-sensitive lipase (Hsl) catalyzes the cleavage of a fatty acyl chain from DG, leaving monoacylglycerol (MG). MG lipases can cleave the last fatty acid freeing the glycerol backbone (Lass et al., 2011). The primary regulatory proteins mediating LD lipolysis are the perilipin (PLIN) proteins (Jackson, 2019). A primary PLIN function is to regulate lipase access to the LD surface. PLIN activity is often modified by targeted phosphorylation. To stimulate lipolysis, PLINs also enhance recruitment of cytoplasmic lipases to the LD surface (Ducharme and Bickel, 2008; Itabe et al., 2017).

*Drosophila* homologues of human Peroxin (PEX) proteins show conserved cellular localization and activity (Anderson-Baron and Simmonds, 2019; Pridie et al., 2020). *Drosophila* and Schneider 2 (S2) cells have also been used extensively to characterize LDs (Beller et al., 2006; Kuhnlein, 2011; Lee et al., 2013) *Drosophila* larval development is strongly influenced by lipid metabolism and the larval adipose tissue localizes to the fat body, which comprises a significant portion of the entire animal (Musselman and Kuhnlein, 2018). *Drosophila* S2 cells can be induced to form LDs reproducibly and are used extensively to study LDs (Beller et al., 2006; Guo et al., 2008; Kory et al., 2015; Krahmer et al., 2011; Sui et al., 2018; Wang et al., 2016; Wilfling et al., 2014; Wilfling et al., 2013). *Drosophila* has only two PLIN homologues: *Lsd-1* and *Lsd-2* (Beller et al., 2006; Bi et al., 2012; Guo et al., 2008). Upon phosphorylation, Lsd-1 facilitates lipid mobilization by recruiting *Drosophila* Hsl to the LD surface, facilitating lipolysis (Bi et al., 2012). Lsd-2 serves to protect the surface of LDs from lipases, such as Brummer (Bmm, (Bi et al., 2012). Bmm is the *Drosophila* ATGL homologue (Gronke et al., 2005).

In yeast cells grown in excess lipid, peroxisomes stably adhere to the surface of LDs (Binns et al., 2006). Subcellular fractionation of yeast cells has identified LDs enriched in peroxisomal β-oxidation enzymes (Binns et al., 2006). Direct peroxisome-LD interaction was observed by TEM in COS7 cells, with clusters of mature peroxisomes adjacent to the surface of LDs (Schrader, 2001). Recently it was shown that the M1 form of Spastin protein interacts with peroxisome resident ATP Binding Cassette Subfamily D Member 1 (ABCD1) to promote interaction of peroxisomes with LDs (Chang et al., 2019). In addition, Pex1, Pex6, and Pex26 were found to be enriched on liver LDs isolated from fasted mice (Kramer et al., 2018). Finally, *C. elegans* PEX5 mediates ATGL translocation to LDs facilitating fasting-induced lipolysis of TGs stored in LDs (Kong et al., 2020). These molecular and physical connections, including sharing of proteins between peroxisomes and LDs strongly suggest regulated trafficking of regulatory proteins between these organelles drives functional coordination.

Formation of LDs in the larval fat body is a tightly regulated process (Kuhnlein, 2012). To probe the role of peroxisomes in tissues where LDs are prominent, we performed a screen for reduced activity of each *Drosophila Pex* gene in the larval fat body. Only a few *Pex* genes showed a fat-body associated phenotype when knocked down by RNA interference (RNAi). The strongest phenotypes were observed when *Pex14* was inhibited, with loss of LDs and reduced survival when animals were fed a high-fat diet. To understand the potential mechanism for a Pex-protein/peroxisomes in LD regulation, we used *Drosophila* S2 cells cultured with excess oleate. Notably, only small differences in peroxisomes were observed when S2 cells were induced to form new LDs. RNA-seq of these cells indicated that a small subset of genes had expression changes. Of the peroxisome-associated genes, only *Pex14* had a significant change in expression. Further, analysis showed that three Pex proteins, Pex3, Pex13, and Pex14 show a high degree of localization to LDs in oleate cultured cells, and this localization occurs independently of other markers indicative of peroxisomes. This recruitment of Pex14 to LDs directly affects lipid storage and subsequent lipolysis as altering the cellular levels of Pex14 by overexpression or RNAi knockdown changed the relative presence of PLIN and lipase proteins at the LD surface.

## Results

### RNAi of Pex genes in the Drosophila fat body differentially affects lipid storage

The fat body comprises a significant proportion of the larval body and is a major contributor to systemic lipid homeostasis (Church and Robertson, 1966; Musselman and Kuhnlein, 2018). During early larval development the fat body enlarges, but in the late stages of development larvae stop feeding and metabolism depends on fat body TG stores (Musselman and Kuhnlein, 2018). Mutations affecting fat storage can be assayed using a buoyancy assay (Reis et al., 2010). To screen for the role of peroxisomes in LD formation we performed a systematic RNAi screen, knocking down each *Pex* mRNA in fat body (Figure 1 A-B, Supplementary Figure 1). RNAi knockdown of most *Pex* genes in the fat body caused some effect on total fat storage, *Pex3*, *Pex14*, and *Pex16* RNAi had the greatest effects (Figure 1C). Closer examination of one *Pex14* RNAi knockdown line (GD2759) confirmed a highly significant reduction in decreased buoyancy (Figure 1D), correlated with reduction in total glycerol content (an indirect measure of TG, 25%, p < 0.01) compared to control larvae (Figure 1E). In wild type fat body, large LD occupied fat body cells (Figure 1F), whereas when *Pex14* RNAi was targeted to the fat body smaller LDs were observed (Figure 1G). The average reduction in LD volume was 11.96μm^3^ (Figure 1H).

**Figure 1.**
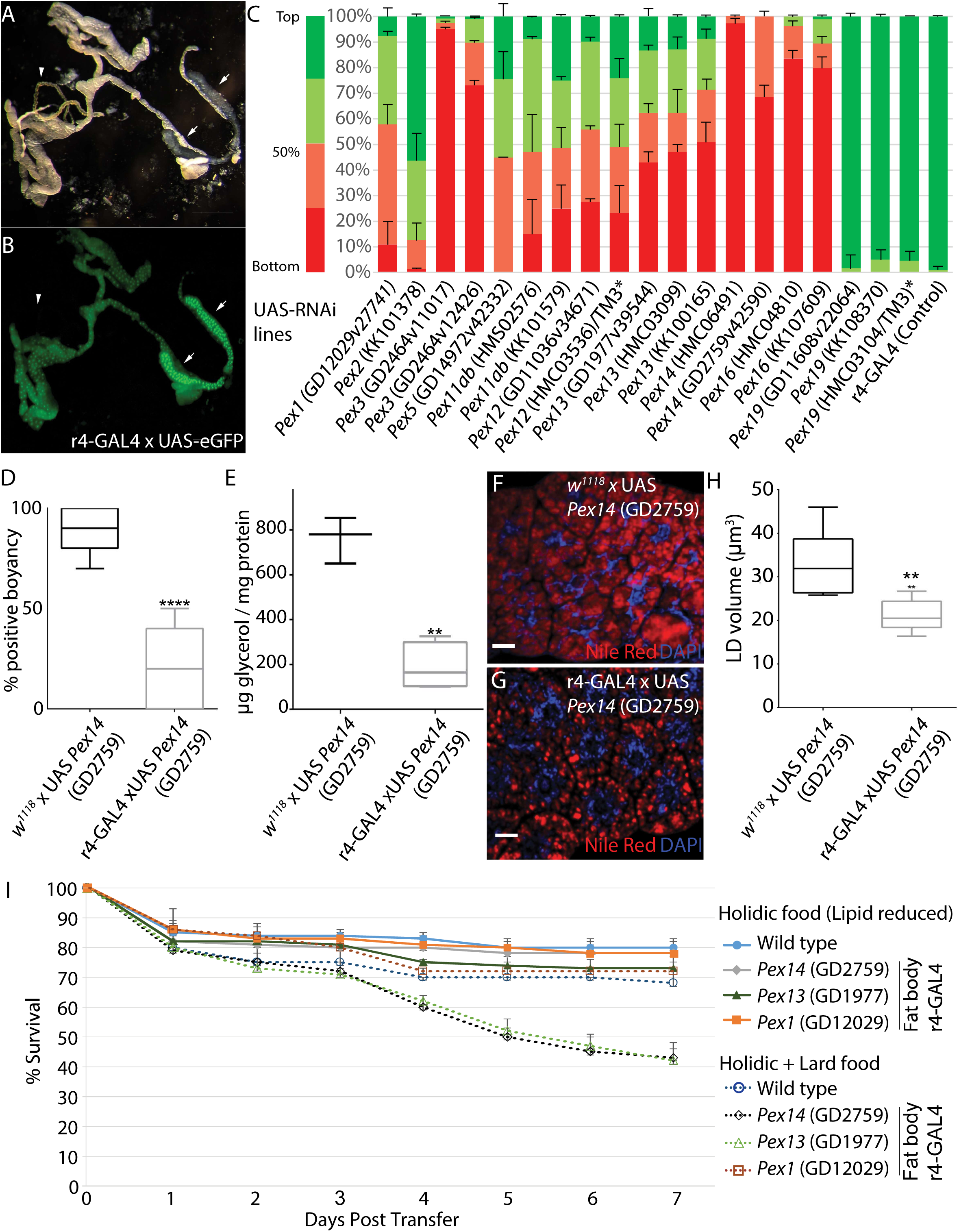
*Pex* gene RNAi knockdown in the fat body affected lipid storge and survival. A) Fat body adhered to salivary glands (arrows) along with Malpighian tubule (arrowhead) dissected from r 4-GAL4xUAS-stinger GFP larvae. B) GFP expression confirmed that the r4-GAL4 driver was active in fat body and salivary glands (arrows). Malpighian tubules (arrowhead) did not express GFP. C) Relative larval buoyancy in 12% sucrose was used as a screen for changes in overall fat-storage. RNAi knockdown screen (r4-GAL4) of most *Pex* genes (*Pex1*, *Pex2*, *Pex5*, *Pex11a/b* and *Pex12*) showed weak buoyancy reduction of (<40% of larva below the top quartile, dark green). *Pex13, Pex14* and *Pex16* crosses had larvae with over 40% in the bottom quartile (dark red). Asterisk indicates heterozygous strains where only homozygous mutant animals were tested. The strongest effects were seen with UAS-Pex14 RNAi (HMC06491 and GD2759). D) Follow up characterization of r4*-GAL4xUAS-Pex14* RNAi (GD2759) animals showed significantly reduced buoyancy compared to a *w^1118^* (no GAL4) control cross (****, p < 0.0001). E) r4*-GAL4xUAS-Pex14* RNAi (GD2759) showed reduced glycerol levels compared to wild type control indicating reduced TG levels (**, p < 0.01). F) Fat body dissected from control animals have multiple large droplets of neutral lipids as evidenced by Nile Red staining in individual cells demarked by nuclear DAPI staining G) r4*-GAL4xUAS-Pex14* RNAi (GD2759) animals have smaller LDs in fat body. Scale bar = 10μm. H) The average volume of fat body LDs is reduced by *Pex14* RNAi (DG2759) (**, p < 0.01). I) Survival of third instar transferred to holidic (lipid reduced) food versus holidic food supplemented with lard. While low fat diet does not significantly affect survival, *Pex13* or Pex14 *RNAi* knockdown in the fat body showed reduced survival compared to *Pex1* RNAi or wild type control animals.

Lipids can be synthesised *de novo* or absorbed from the diet by the gut. *Drosophila* larvae store TG in LDs within the fat body to fuel subsequent pupal development (Heier and Kühnlein, 2018). Third instar larvae were raised on a chemically defined (holidic, (Piper et al., 2014) diet where the only added lipid was cholesterol as *Drosophila* are cholesterol auxotrophs. (Vinci et al., 2008). Larvae with *Pex13*, *Pex14,* or *Pex1* RNAi knockdown in the fat body (Figure 1I) survive equally well on a holidic diet. Larvae will consume a lipid rich diet when lard is added to their food (Woodcock et al., 2015). When flies were raised on holidic + lard food, the survival rate of wild type or *Pex1* RNAi fat body knockdown larvae was like those raised on holidic food. However, fat body RNAi knockdown of *Pex13* or *Pex14* strongly reduced survival on lard-supplemented food (Figure 1I).

### RNA-Seq of S2 cells in conditions promoting LD formation or lipolysis

*Drosophila* S2 cells rapidly form LDs that are consistent in both number and volume (Guo et al., 2008) when cultured medium supplemented with oleate (+Oleate) (Darfler, 1990), an 18-carbon monounsaturated fatty acid. +Oleate culture conditions caused relatively little change in peroxisome number (Figure 2A). To determine the differential regulation of peroxisome versus LDs, RNA sequencing was used to compare cells cultured in Schneider’s medium + FBS (Standard) versus +Oleate conditions. This identified 249 mRNAs that consistently showed significant changes in relative abundance (n=3, padj<0.1, Figure 2B, Supplementary Table 1). Gene Ontology (GO) clustering showed differentially expressed mRNAs encoded proteins linked to mitochondria, peroxisomes, and the endomembrane system (Supplementary Table 2). In terms of predicted molecular function, there was enrichment of mRNAs encoding multiple proteins involved in peroxisomal fatty acid β-oxidation (Supplementary Table 2). However, this was not paired with a corresponding increase in mRNAs encoding the peroxisome biogenesis (Pex) factors required for peroxisome proliferation. In fact, only one *Pex* mRNA, *Pex14*, was significantly (padj < 0.1) enriched in S2 cells cultured in +Oleate conditions (Supplementary Table 1). Increased expression of *Pex14*, and a slight increase in *Pex13* was detected by quantitative RT-PCR (qRTPCR) in +Oleate cultured S2 compared to Standard conditions (Figure 2C). Relatively less change was observed other *Pex* genes (Supplementary Table 1), *e.g., Pex2* (Figure 2C). When cells were transferred to Lipolytic conditions, levels of *Pex2*, *Pex13*, and *Pex14* mRNA were all elevated compared to standard culture, but to a lesser extent (Figure 2C).

**Figure 2.**
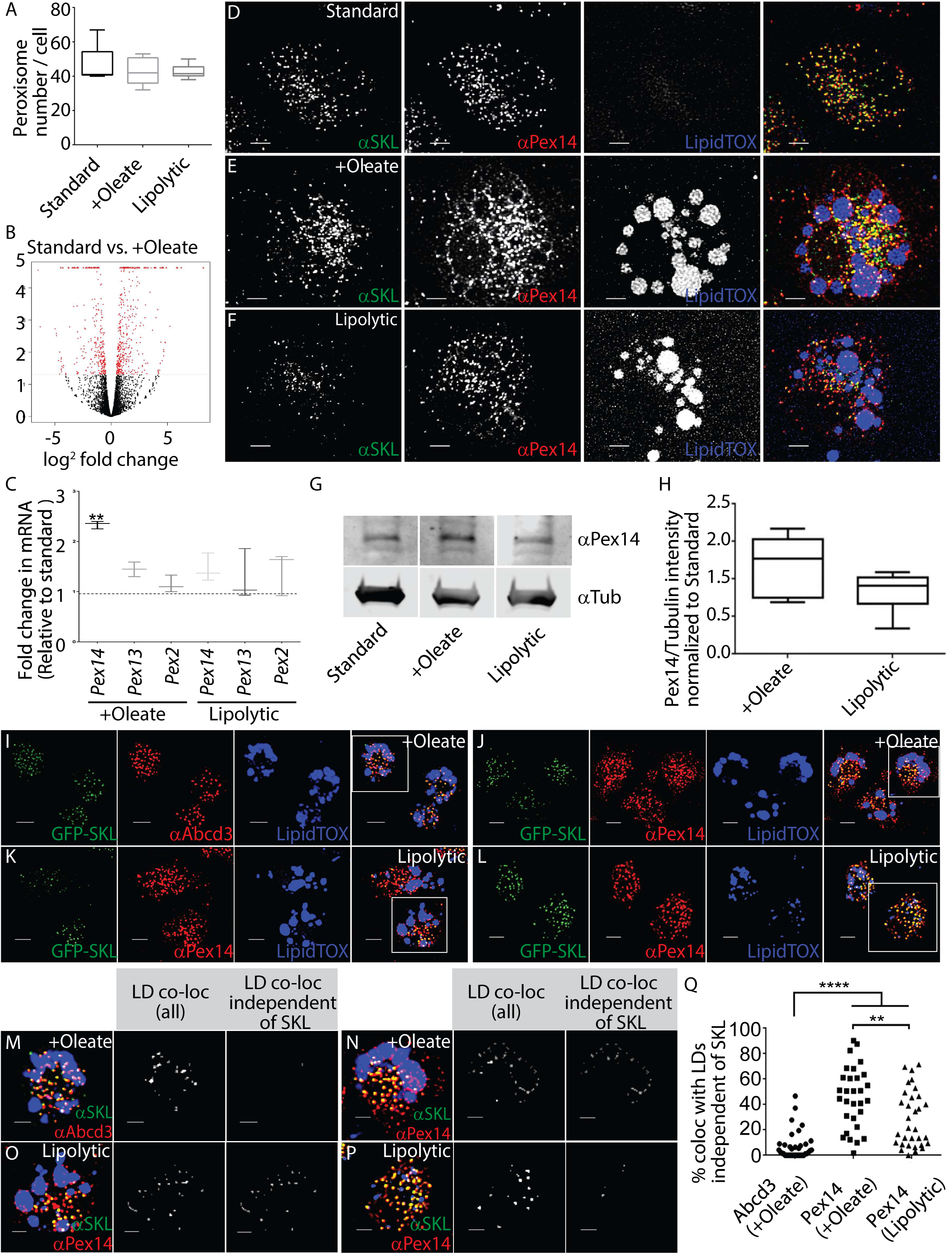
Culture of S2 cells in conditions promoting LD formation or lipolysis caused changes in *Pex* gene expression and Pex protein localization. A) The average number of peroxisomes observed in cells cultured in Standard, +Oleate, or Lipolytic culture conditions. B) A volcano plot comparing differences in mRNA expression in S2 cells in Standard versus +Oleate conditions. Red dots represent those mRNAs with statistically significant differences between the two conditions. Complete mRNA SEQ data is provided as Supplementary Tables 1-2. C) Quantitative RT-PCR can detect altered levels of *Pex14* relative to *Pex2*, *Pex3* or *Pex13* mRNAs in S2 cells in +Oleate or Lipolytic conditions (**, p < 0.01). D) In S2 cells cultured under Standard conditions, markers for mature peroxisomes (green, αSKL) and endogenous Pex14 (red, αPex14) signal are overlapping (yellow), and few LDs are present (blue, LipidTOX). E) When S2 cells were cultured in +Oleate conditions a portion of the Pex14 signal that does not overlap with the mature peroxisome marker SKL was observed surrounding LDs. F) When cells are transferred from +Oleate to Lipolytic conditions, LDs were more numerous and smaller (indicating fragmentation) and the SKL-independent Pex14 signal is seen only surrounding larger LDs. Scale bars = 2nm. G) A representative western blot showing relative levels of Pex14 (αPex14) normalized to tubulin (αTub) in cells cultured in Standard, +Oleate and Lipolytic conditions. H) Quantification of expression of Pex14 relative to Tubulin in cells cultured in +Oleate and Lipolytic conditions, relative to Standard conditions (n=3). I) Peroxisome markers GFP-SKL (green, matrix) and αAbcd3 (red, membrane) are colocalized in +Oleate cells. Boxed region is showed magnified in M. J) In +Oleate cells αPex14 (red) are seen, one overlapping with GFP-SKL and the other co-localizing with the edge of LDs (blue, LipidTOX). Boxed region showed is magnified in N. Two distinct populations of cells were observed when cells are transferred from +Oleate to Lipolytic culture conditions. K) In approximately half the cells, a pattern of Pex14 localization like +Oleate conditions is seen. Boxed region showed is magnified in O. L) In the other half of the cells, the LDs (blue, LipidTOX) are fragmented and Pex14 is not associated with the perimeter of LDs. Boxed region showed is magnified in P. Scale bars = 10nm. M-P) Zoomed images of examples cells used for quantification of proportion of signal colocalizing with LDs with and without also colocalizing with SKL. M) Mature peroxisome marker, Abd3 (red, αAbcd3) at LDs is largely overlapping with SKL (green, αSKL) in +Oleate cells. N) A large proportion of Pex14, (red, αPex14) associated with LDs (blue, LipidTOX) is not colocalized with SKL in +Oleate cells. O) When cells are transferred to Lipolytic conditions. approximately half the have a pattern of Pex14 LD co-localization like +Oleate. P) In the other half of the cells LDs (blue) are fragmented and Pex14 associated with LDs becomes strongly co-localized with the mature peroxisome marker SKL. Q) Quantification of the proportion of Abcd3 or Pex14 signal co-localizing with LDs independently of SKL. All images are shown as maximum intensity projections of deconvolved Z stacks.

### Pex14 localized to LDs when S2 cells were cultured in +Oleate conditions

When S2 cells were cultured in standard conditions, the punctate signal from anti-SKL (mature peroxisomes) and anti-Pex14 largely overlapped (Figure 2D). When S2 cells were cultured in +Oleate conditions, additional Pex14 signal surrounded LDs, independently of SKL (Figure 2E). When S2 cells were subsequently transferred to Lipolytic conditions, less punctate Pex14 independent of SKL signal was observed, except surrounding the large LDs (Figure 2F). Western blotting showed that endogenous Pex14 protein levels were elevated in +Oleate S2 cells but far less so in Lipolytic conditions (Figure 2G-H). In + Oleate cells, peroxisome marker protein membrane-associated ATP binding cassette subfamily D member 3 (Abcd3, also known as Pmp70) did not localize to LDs except when part of peroxisomes as evidenced by co-localization with SKL (Figure 2 I), whereas much of the Pex14 signal was independent of SKL (Figure 2J). When cells were transferred to Lipolytic conditions, two phenotypes were observed, when large LDs were present, they were surrounded by peroxisome independent Pex14 (Figure 2K), whereas when only small LDs were present, Pex14 largely overlapped with peroxisomes (Figure 2L). Quantification of the three-dimensional co-localization of individual cells of each category showed that Pex14 co-localization to LDs independently of SKL was significantly higher than Abcd3 (Figure 2M-Q).

### Pex14 RNAi knockdown altered TG lipolysis and LD morphology

In S2 cells treated with *Pex14* dsRNA, PTS1-mediated (SKL) peroxisomal import was shown previously to be reduced (Mast et al., 2011). No change in the spatial distribution of peroxisomes relative to LDs was detected in +Oleate or Lipolytic cultured cells treated with *Pex14* dsRNA (Figure 3A-D). In +Oleate cells *Pex14* RNAi treatment significantly reduced the average volume of peroxisomes relative to control cells (Figure 3E). When cells were transferred to Lipolytic conditions this relative difference was not longer present (Figure 3E). Conversely *Pex14* RNAi had no significant effect on peroxisome number in +Oleate cells but did strongly suppress peroxisome number when cells were later (+48h) transferred to Lipolytic conditions (Figure 3F). Increased LD number and volume in +Oleate conditions has been shown previously to be a function of increased TG storage in S2 cells (Guo et al., 2008). *Pex14* dsRNA did not significantly affect the rate of TG storage in LDs in +Oleate cells (Figure 3G-H). However, *Pex14* RNAi had a significant effect on LD volume and number when cells were later (+48h) transferred to Lipolytic conditions (Figure 3G-H). Colorimetric assays measuring lipoprotein lipase-induced changes in glycerol levels have been shown previously to largely correspond to TG levels in *Drosophila* cell lysates (Tennessen et al., 2014). *Pex14* RNAi treatment had a significant effect on TG levels when cells were transferred from +Oleate to Lipolytic conditions (Figure 3I). This suggested Pex14 regulation of lipolysis of TGs that was separable from its function at peroxisomes. Reduction in *Pex14* mRNA by RNAi treatment was confirmed by qRTPCR in all experiments (Figure 2J).

**Figure 3.**
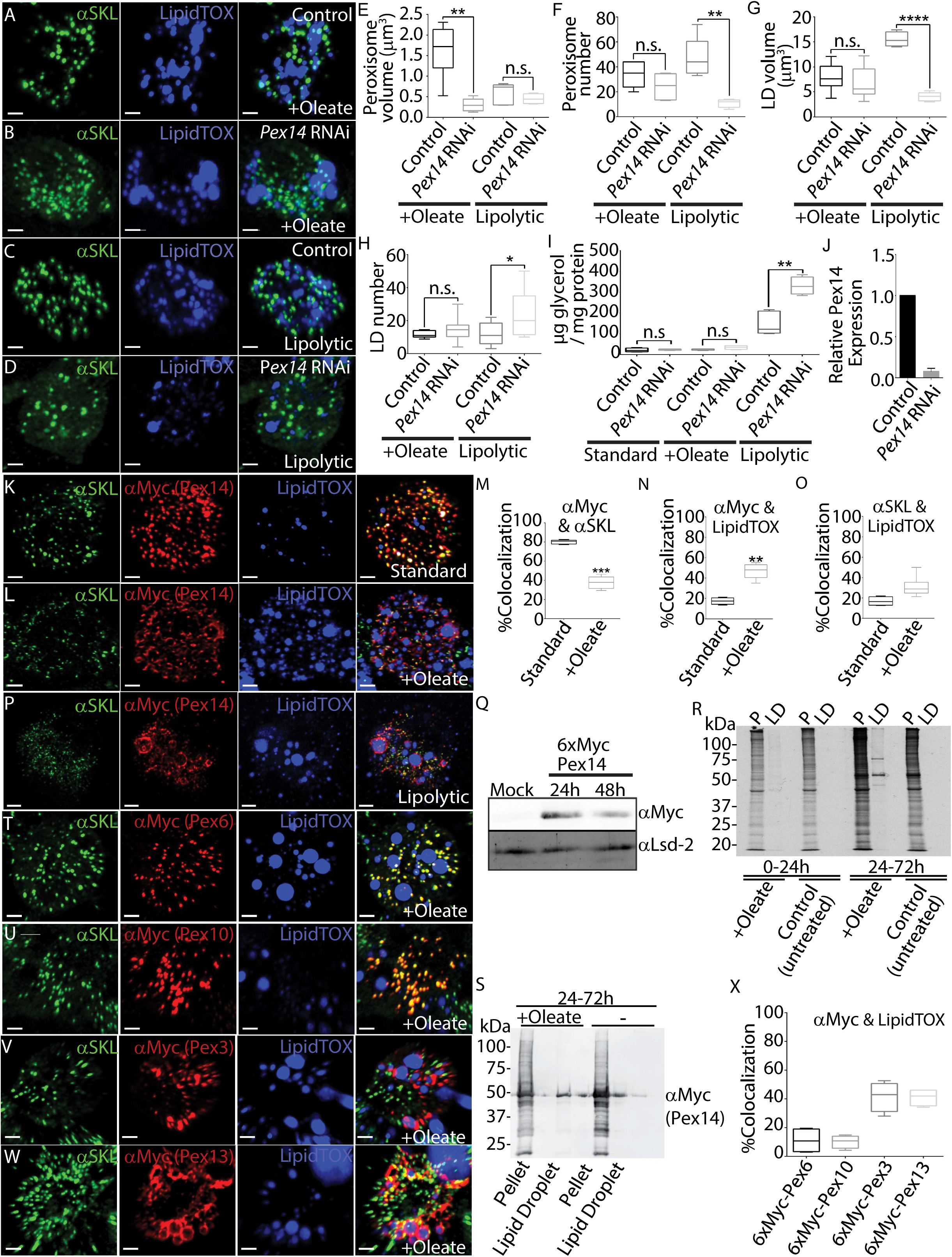
Pex14 and two other Pex proteins, Pex3 and Pex13, localized to LDs. A) S2 cells cultured in +Oleate conditions treated with scrambled dsRNA (Control). B) S2 cells cultured in +Oleate conditions treated *Pex14* dsRNA (*Pex14* RNAi). Punctate anti-SKL signal marks peroxisomes (green, αSKL), and LipidTOX Deep Red marks LDs (blue, LipidTOX). C) S2 cells in Lipolytic culture conditions treated with scrambled dsRNA (Control) or D) S2 cells in Lipolytic culture conditions treated with *Pex14* dsRNA (*Pex14* RNAi). Scale bar = 2μm. E) The average volume of peroxisomes per S2 cell cultured under +Oleate or Lipolytic conditions treated with *Pex14* dsRNA (*Pex14* RNAi) relative to control (**, p < 0.01). F) The average number of peroxisomes per S2 cell cultured under +Oleate or Lipolytic conditions treated with *Pex14* dsRNA (*Pex14* RNAi) relative to control (**, p < 0.01) G) The average volume of LDs per cell when cultured under +Oleate or Lipolytic conditions treated *Pex14* dsRNA (*Pex14* RNAi) relative to control (****, p < 0.0001). H) The average number of LDs per cell when cultured under +Oleate or Lipolytic conditions treated *Pex14* dsRNA (*Pex14* RNAi) relative to control *, p < 0.05). I) The amount of free glycerol present in the medium of S2 cells cultured in Standard, +Oleate and Lipolytic conditions treated with a scrambled dsRNA amplicon or dsRNA targeting Pex14 (RNAi, **, p < 0.01). J) Levels of *Pex14* mRNA measured by qRTPCR from S2 cells treated with scrambled (control) or Pex14 dsRNA (RNAi) confirm knockdown efficiency. K) S2 cells transfected with 6xMyc-Pex14 (red, αMyc) and cultured in Standard culture conditions. Punctate SKL (green αSKL) marks peroxisomes L) In S2 cells expressing 6xMyc-Pex14 cultured in +Oleate conditions 6xMyc-Pex14 signal was seen surrounding a subset of LDs independently of SKL. Each image is a maximum projection of a three-dimensional volume encompassing the entire cell. Scale bar = 2μm. M) Quantitation of colocalization between 6xMyc-Pex14 and mature peroxisomes (αSKL). N) Quantitation of colocalization between 6xMyc-Pex14 and LDs. O) Quantitation of colocalization between mature peroxisomes (αSKL). and LDs (**, p < 0.01. ***, p < 0.001). P) Like what was observed with endogenous Pex14, in S2 cells in Lipolytic conditions Myc-Pex14 surrounds only large LDs. Q) Western blotting shows that 6xMyc-Pex14 is part of the LD fraction (also contains Lsd-2) of cells cultured +Oleate conditions for 24h or 48h probed with anti-Myc (Pex14) and anti-Lsd-2 R) ^35^S-Met pulse-chase labelling shows that newly synthesized 6xMyc-Pex14 is recruited to LDs. Cells were incubated with ^35^S-Met immediately after transfection (0-24h) or 24h after transfection (24-72h). Cells were cultured in +Oleate conditions or in control (untreated) conditions. Several radiolabelled bands were observed indicating that newly synthesized proteins are recruited to LDs that form in +oleate cells (24-72h). S) A western blot of the 24-72h LD fractions shows that 6xMyc-tagged Pex14 (αMyc) is present at LDs at much higher levels in +Oleate cultured cells. T-W) S2 cells expressing 6xMyc tagged (T) Pex6, (U) Pex10, (V) Pex3 or (W) Pex13 cultured for 48h in +Oleate conditions. Punctate anti-SKL (green, αSKL) marks mature peroxisomes, anti-Myc (red, αMyc) marks tagged proteins (red), LipidTOX Deep Red marks LDs (blue). Scale bar = 2μm. X) Quantitation of colocalization between the Myc and LD (LipidTOX) signals shows that Pex3 and Pex13 but not Pex6 or Pex10 are strongly recruited to LDs like Pex14 in +Oleate cultured cells. Images are shown as maximum projections of a three-dimensional volume encompassing the entire cell.

In metazoan cells, peroxisomes are continually regenerated with an approximate half-life of approximately 2 days, in a process that requires 12 conserved Pex proteins (Nordgren et al., 2013). Myc-tagged *Drosophila* Pex proteins, except Pex3, largely overlap with the SKL peroxisome marker in S2 cells (Baron et al., 2016) including Pex14 (Figure 3K). Myc-tagging did not affect peroxisome (SKL) independent localization of Pex14 to LDs in +Oleate cultured cells (Figure 3L-O). Pex14 is a highly stable protein with a relatively low rate of turnover (Natsuyama et al., 2013). When S2 cells were cultured in transferred (+48h) from +Oleate to Lipolytic conditions 6xMyc-Pex14 remained associated with LDs independently of mature peroxisomes (Figure 3P). 6xMyc-Pex14 was present in the protein fraction of LDs isolated +Oleate cells for at least 48h (Figure 3Q). Pulse-chase radioactive protein labelling showed multiple newly synthesised proteins are recruited to LDs in +Oleate cells (Figure 3R). Western blotting of the fractions showed that the proportion of 6xMyc-Pex14 in the LD fraction was elevated when cells were cultured in +Oleate conditions for 24-72hr (Figure 3S).

### Pex3, Pex13 are Pex14 are the only Pex proteins localized to LDs independently of peroxisomes

We showed previously that except for Pex3, Pex7 and Pex19, Myc-tagged *Drosophila* Pex proteins are localized to mature (SKL-important competent) peroxisomes (Baron et al., 2016). A similar recruitment specifically to peroxisomes is seen in +Oleate cells (Figure 3T-U). However, like Pex14, Pex3 and Pex13 localized to LDs independently of peroxisomes (marked by SKL) when cells were cultured in +Oleate conditions (Figure 3V-X).

### Pex14 localizes to LDs in a Pex19 independent manner

Pex19 is predicted to be required for membrane insertion of Pex3, Pex13 and Pex14 (Itoh and Fujiki, 2006). In +Oleate S2 cells, a significant portion of the Pex14 can be observed surrounding a subset of LDs while the rest were localized to peroxisomes (Figure 2Q). Pex14 was shown previously to be localized to mitochondria in human fibroblasts when Pex19 is absent (Sacksteder et al., 2000). In +Oleate S2 cells, Pex14 surrounding LDs was independent from a mitochondrial marker, Cytochrome C (CytC, Figure 4A). S2 cells deleted for *Pex19* (Pex19KO) that were also expressing neonGreen-SKL, no punctate signal was observed corresponding to mature peroxisomes (Figure 4B). However, in these same cells, Pex14 still surrounded LDs in +Oleate conditions (Figure 4B). CytC and SKL signal does overlap in some places in the cytosol, but not adjacent to LDs (Figure 4C). Pex16 is also needed for PMP insertion, and in *Pex16* RNAi knockdown cells (+Oleate) Pex14 does not localize to peroxisomes or LDs (Figure 4D). SKL, Pex14 and CytC did not appreciably co-localize in Pex19KO cells under standard culture conditions (Figure 4E). In Pex19KO cells (+Oleate) Pex14 was largely unassociated with CytC but very strongly localized to LDs (Figure 4F).

**Figure 4.**
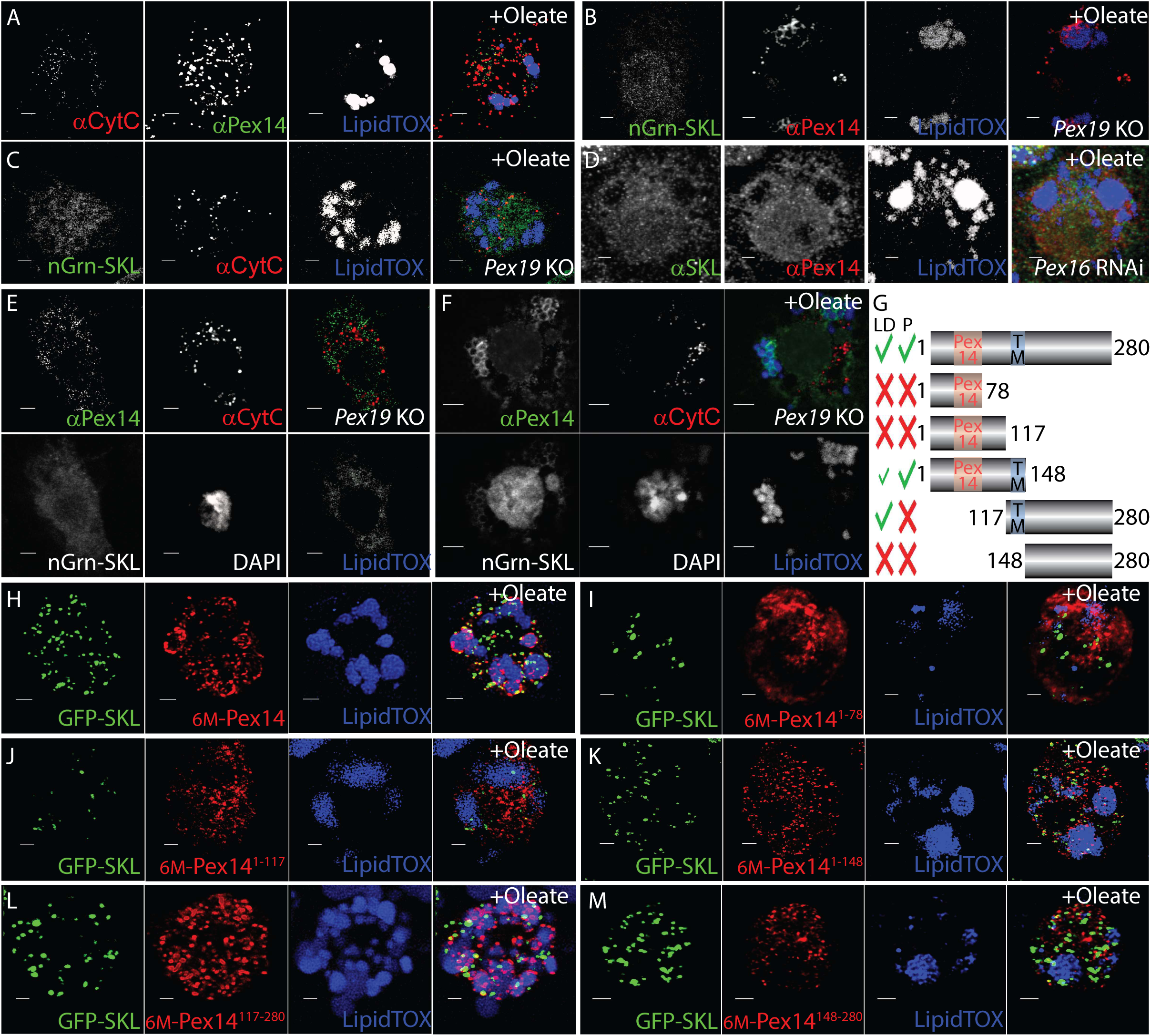
The C-terminal region of Pex14 mediates LD association. A) Endogenous Pex14 (green, αPex14) does not co-localize with mitochondrial marker cytochrome c (red, αCytC) in +Oleate cells. B) In *Pex19*KO cells, Pex14 (red, αPex14) is localized only to LDs and does not co-localize with neonGreen-SKL (green, mgrn-SKL). C) In *Pex19*KO cells, mitochondria (red, αCytC) do co-localize with neonGreen-SKLSKL (green, nGrn-SKL). D) RNAi knockdown of Pex16 suppresses localization of Pex14 (red, αPex14), peroxisome formation (no punctate SKL, green, αSKL) or with LDs (blue, LipidTOX). E) Pex14 (green, αPex14) and cytochrome c (red, αCytC) visualized simultaneously with neonGreen -SKL and LDs (blue, LipidTOX) in *Pex19*KO cells cultured in Standard conditions. F) When *Pex19*KO cells are transferred +Oleate conditions Pex14 (green, αPex14) surrounds LDs. G) *Drosophila Pex14* encodes a single protein isoform that is 280 amino acids long and contains a conserved Pex14 domain (red) with a single internal transmembrane (TM) domain (blue). Myc-tagged transgenes expressing truncations containing the N-terminal Pex14/TM domain (Pex14^1-148^), the C-terminal region plus TM domain (Pex14^117-280^) and the C-terminal domain alone (Pex14^148-280^). Truncations containing the Pex14 region localized to peroxisomes (P), while all constructs containing the TM domain localized to LDs. H) In +Oleate cultured cells, full length 6xMyc tagged Pex14 (red) co-localized peroxisomes (punctate green GFP-SKL) and to LDs (blue, LipidTOX). I-J) Neither Pex14^1-78^nor Pex14^1-117^ o-localized with peroxisomes or LDs. K) Pex14^1-148^ co-localized with peroxisomes, partially co-localized with LDs. L) Pex14^117-280^ formed a punctate pattern that was localized almost exclusively to the periphery of LDs. M) Pex14^148-280^ forms a punctate pattern that is distinct from both the LDs and peroxisomes. All images are maximum projections of confocal images encompassing the entire cell volume. Scale bar = 2μm.

### The C-terminal region of Pex14 was required for LD association

In 2019, Barros-Barbosa et al. showed that rat PEX14 is an intrinsic membrane protein with an N-in C-out topology (Barros-Barbosa et al., 2019; Reuter et al., 2021). Myc-tagged N-and C-terminal truncations of Pex14 were tested for localization to LDs or peroxisomes in +Oleate cells (Figure 4G). The N-terminal 78 amino acid region of Pex14 is homologous to a domain in mammalian PEX14 that associates strongly with microtubules (Bharti et al., 2011) *Drosophila* Pex14 amino acids 1-148 include a region homologous to the N-terminal ‘Pex14’ domain in yeast and human homologues (Mast et al., 2011). This region also contains a predicted transmembrane domain (TM, aa 117-148 Figure 4G), homologous to yeast Pex14p or mammalian PEX14 (Niederhoff et al., 2005; Oliveira et al., 2002). Full length 6xMyc-Pex14 localized to LDs in +Oleate (+48h) S2 cells (Figure 4H). Pex14^1-78^ did not co-localize with LDs or peroxisomes (Figure 4I). The N-terminal Pex14^1-117^ (lacking the TM domain) also did not co-localize with LDs or peroxisomes (Figure 4J). However, Pex14^1-148^, which encompassed the N-terminal half of Pex14 including the TM domain co-localized with GFP-SKL (peroxisomes) but not LDs (Figure 4K). The C-terminal half of Pex14 that includes the TM domain (Pex14^117-280^) localized to LDs but not peroxisomes (Figure 4L). Pex14^148-280^ encompassing the C-terminal half of Pex14 but lacks the TM domain localized to neither peroxisomes nor LDs (Figure 4M).

### Directly altering the amount of Pex14 at the LD surface affected recruitment of lipases

The mRNA encoding *Drosophila Hormone sensitive lipase (Hsl)* was also relatively much higher in S2 cells cultured in +Oleate conditions (Supplementary Table 1). Thus, we examined the relative recruitment of LD lipases Bmm and Hsl to LDs in S2 cells where the level of Pex14 was elevated via transgene expression. When S2 cells were transferred Lipolytic culture conditions for 24 h, 3xFLAG-Bmm surrounded the LD periphery, especially those that were smaller than when cells were in +Oleate conditions (Figure 5A). When Lipolytic cells were co-transfected with *Bmm* and *Pex14* transgenes both Bmm and Pex14 co-localized at the LD surface (Figure 5B). The DG lipase HSL also surrounded LDs in Lipolytic S2 cells (Figure 5C). When *Hsl* and *Pex14* were co-overexpressed, Hsl was observed in a cytosolic punctate pattern distinct from LDs, while Pex14 surrounded a subset of relatively large LDs (Figure 5D). 50-60% of the total signal from Myc-Pex14 was recruited to LDs even if Bmm or Hsl levels were also elevated (Figure 5E). However, colocalization between FLAG-Hsl and LDs (LipidTOX) was reduced significantly from 62.2% to 35.9% when Myc-Pex14 was co-overexpressed (Figure 5F). LD volume and number per cell were relatively unaffected when FLAG-Bmm and Myc-Pex14 were co-overexpressed (Figure 5G-H). However, expressing FLAG-Hsl and Myc-Pex14 simultaneously caused a significant decreased LD number and increased LD volume (Figure 5G-H). Increased number and decreased LD volume indicates elevated lipase activity (Marcinkiewicz et al., 2006).

**Figure 5.**
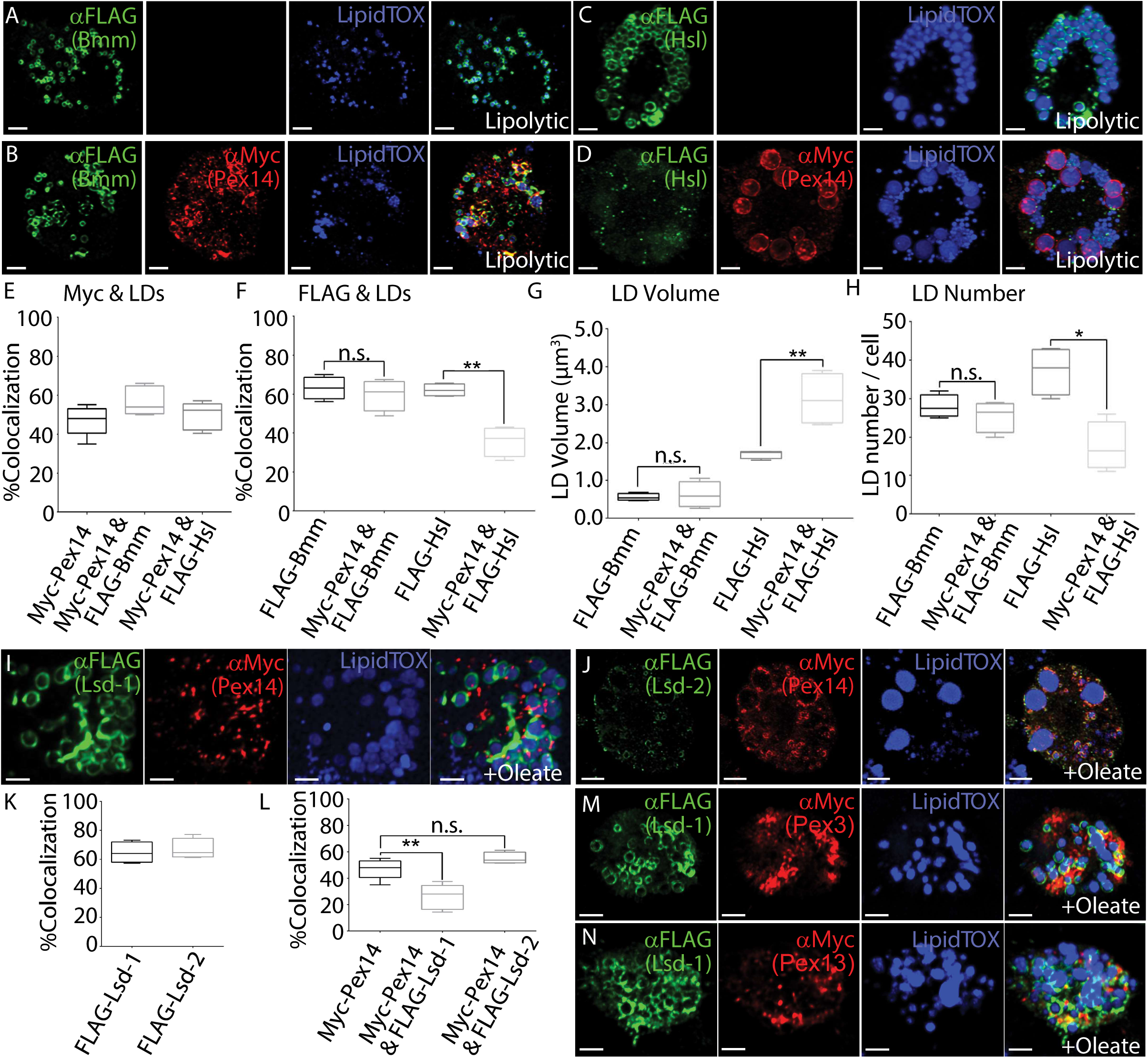
Pex14 affects lipid storage through Hsl recruitment to the LD surface. A) 3xFLAG-Bmm (green, αFLAG) overexpressed in Lipolytic S2 cells surrounded small LDs (blue, LipidTOX Deep Red). B) In Lipolytic cells co-overexpressing 6xMyc-Pex14 (red, αMyc) and 3xFLAG-Bmm (green, αFLAG), Bmm localization to LDs was unaffected. C) When 3xFLAG-Hsl (green, αFLAG) is overexpressed in Lipolytic S2 cells it surrounds LDs (blue, LipidTOX Deep Red). D) In Lipolytic cells co-overexpressing 6xMyc-Pex14 (red, αMyc) and 3xFLAG-Hsl (green, αFLAG), Hsl was excluded from the large LDs, especially those where Pex14 is present. E) Quantification of colocalization showed that Pex14 recruitment to LDs was largely unaffected by increased levels of Bmm or Hsl. F) Quantitative analysis of Lipolytic cells co-overexpressing Myc-Pex14 showed no effect on Bmm recruitment to LDs. When Myc-Pex14 is co-overexpressed, Hsl surrounding LDs was reduced by ∼35% (**, p < 0.01). G) In Lipolytic cells, co-overexpressing FLAG-Bmm and Myc-Pex14 shows no change in LD volume. Overexpressing FLAG-Hsl and Myc-Pex14, LD increased average size by ∼1.7x (**, p < 0.01). H) In Lipolytic cells overexpressing FLAG-Bmm and Myc-Pex14 LD number was unaffected. In cells overexpressing FLAG-Hsl and Myc-Pex14, LD number decreased ∼2-fold. (*, p < 0.04). H) In +Oleate cells co-overexpressing 6xMyc-Pex14 and 3xFLAG-Lsd-1, Pex14 does not localize to LDs. J) In +Oleate cells co-overexpressing 6xMyc-Pex14 and 3xFLAG-Lsd-2, Myc-Pex14 is recruited to all LDs. K) Overexpressed 3xFLAG-tagged Lsd-1 and Lsd-2 were both strongly recruited (>60%) to LDs in +Oleate culture conditions. L) Overexpression of Lsd-1 suppresses (- 1.8x) recruitment of Pex14 to LDs (**, p < 0.01). M) +Oleate S2 cells co-overexpressing 3xFLAG-Lsd-1 6xMyc-Pex3 did not localize to LDs. N) +Oleate S2 cells co-overexpressing 3xFLAG-Lsd-1 6xMyc-Pex13 to did not localize LDs. Each image shows a maximum projection of a three-dimensional volume encompassing the entire cell. Scale bar = 2μm.

### Pex14 recruitment to LDs was affected by Drosophila perilipin proteins

*Drosophila* has two PLIN proteins, Lsd-1 and Lsd-2. While there is some overlap in their activities, generally, Lsd-1 facilities LD lipid mobilization by Hsl while Lsd-2 suppresses Bmm-mediated lipolysis at LD (Beller et al., 2010; Marcinkiewicz et al., 2006). Thus, we examined how altered Lsd-1 or Lsd-2 affects Pex14 recruitment to LDs. In +Oleate S2 cells simultaneously overexpressing tagged Pex14 and Lsd-1, Pex14 was prevented from being localized to LDs (Figure 5I). Conversely, when Lsd-2 and Pex14 were co-overexpressed, both localized to the LD surface (Figure 5J). In +Oleate cultured S2 cells, FLAG-Lsd-1 and FLAG-Lsd-2 showed approximately 70% colocalization with the LD surface (Figure 5K). However, recruitment of Myc-Pex14 to LDs was significantly lower (26.5%, p < 0.01) when FLAg-Lsd-1 was simultaneously overexpressed (Figure 5L). Overexpression of FLAG-Lsd-1 had similar effect on localization of Myc-Pex3 or Myc-Pex13 to LDs (Figure 5M-N).

### Recruitment of Pex14 to LDs is conserved in mammalian NRK and Huh7 cells

When NRK cells were cultured in DMEM + 10% FBS (Standard) conditions, PEX14 was largely recruited to mature peroxisomes marked by ATP-binding cassette sub-family D member 3 (ABCD3, Figure 6A). When NRK cells were cultured in DMEM +10% FBS + 1 mM oleate for 48h (+Oleate), a significant portion of PEX14 was observed as punctate signal surrounding LDs that was distinct from ABCD3 (Figure 6B). A similar effect was seen in +Oleate cultured Huh7 cells (Figure 6C-D). Colocalization between PEX14 and the LD surface increased significantly from 3.5% in Standard conditions to 42.8% in + Oleate conditions (Figure 6E). However, localization of a mature peroxisomes (ABCD3) did not change significantly when NRK cells were cultured in Standard or +Oleate conditions (Figure 6F). Like what was observed in S2 cells, no change in peroxisome volume was observed in +Oleate NRK cells (Figure 6G), however peroxisome number increased significantly (Figure 6 H). Like what occurred in +Oleate cultured S2 cells, PEX14 levels were elevated in NRK cells cultured in +Oleate at 24h and persisting for at least 48h (Figure 6 I-J). TEM-immunolabelling of +Oleate NRK cells showed that PEX14 was associated with membranes, including those of presumptive LDs or LD-associated vesicles ranging in size from 40-100 nm in diameter (Figure 6K-L). The majority of PEX14 was localized within 40nm of the LD surface (Figure 6M).

**Figure 6.**
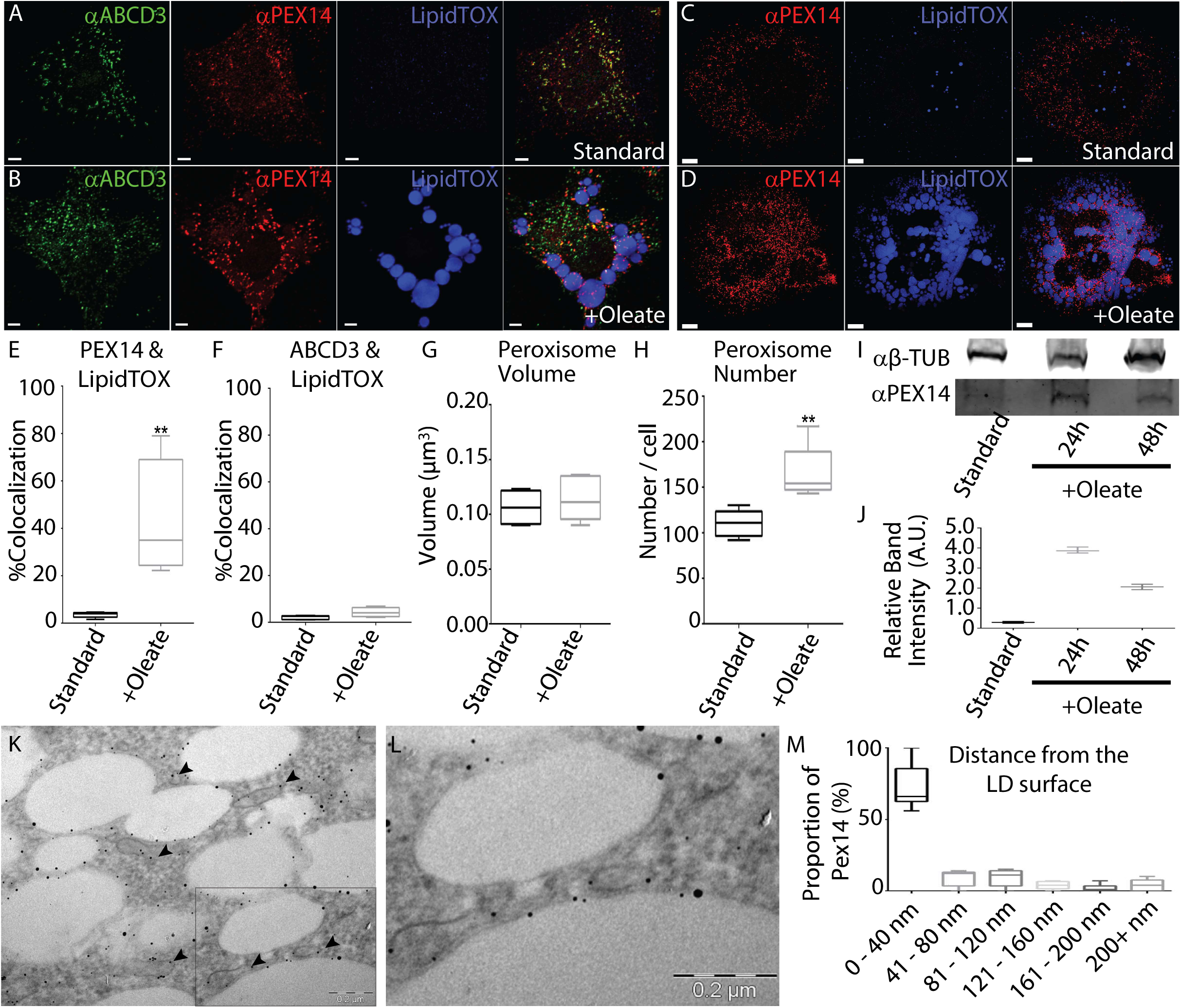
Pex14 localization to LDs is conserved in mammalian cells. A) In NRK cells cultured in Standard conditions, peroxisome marker ATP Binding Cassette Subfamily D Member 3 (green αABCD3) co-localized with PEX14 (red, αPEX14) and few LDs are present (blue, LipidTOX). B) In +Oleate cultured cells, PEX14 was observed surrounding LDs independently of ABCD3. C) When Huh7 cells were cultured Standard conditions PEX14 was not recruited to LDs. D) In + Oleate conditions large LDs formed and were surrounded by PEX14. Each representative image is shown as a maximum projection of a three-dimensional volume encompassing the entire cell. Scale bar = 2μm. E) Colocalization between Pex 14 with LDs (LipidTOX) increased in +Oleate conditions (**p< 0.01). F) Average colocalization between ABCD3 (mature peroxisomes) and LDs is largely unaffected when NRK cells were cultured in +Oleate conditions. G) Average peroxisome volume remains constant in NRK cells cultured under Standard or +Oleate conditions H) The average number of peroxisomes increased when NRK cells were transferred to +Oleate conditions (**, p < 0.01). I) Immunoblotting with anti-PEX14 (αPEX14) showed that levels of PEX14 in NRK +Oleate cell lysates increased at 24 h or 48 h compared to cells cultured in Standard conditions relative to the loading control β-tubulin (αβ-TUB). J) Immunoblot quantification shoed that the relative level of PEX14 increased 4-fold after 24h of +Oleate culture and remained elevated (2-fold) after 48h. K) TEM images of NRK cells probed with anti-PEX14 (black spots) showed PEX14 associated with vesicle-like structures 40-100 nm in diameter (arrowheads) adjacent to LDs. L) A magnified view of the boxed region is shown magnified in (K). M) Quantification of the proportion of PEX14 signal relative to the LD surface. from 10 individual TEM images. Scale bar = 0.5μm.

## Discussion

There is a lipid-responsive localization of a subset Pex proteins (Pex3, Pex13 and Pex14) that cooperate in the early stages of peroxisome proliferation to LDs that occurs independently of markers associated with mature peroxisomes. The primary driver of this activity appears to be Pex14. When associated with LDs, Pex14 affects recruitment of lipases and PLIN proteins and influences storage and mobilization of TGs from LDs. However, the localization to, and regulation of LDs observed was different from that previously reported in terms of *PEX14* influence on lipid metabolism in mammalian cells. Yin *et al*. performed a microarray analysis in human liposarcoma SW782 cells to identify genes upregulated during the late stages of adipogenesis (Yin et al., 2014). Of the 11 mRNAs that showed a greater than 10-fold increase, one was *PEX14* (Yin et al., 2014) This is similar to what was observed in our RNA-SEQ comparison of S2 cells cultured in Standard and +Oleate conditions (Supplementary Tables 1-2, Figure 2C). Other groups similarly found *PEX14* upregulation during adipocyte differentiation in SW872 cells (Zhu et al., 2014). This was assumed to be a prelude to peroxisome proliferation. However, we found that peroxisome number does not increase in +Oleate cells and expression of other Pex genes needed for making new peroxisomes (*e.g.*, *Pex2*) did not change (Supplementary Tables 1-2, Figure 2A-C). This suggests that the LD-associated activity of Pex14 we identified is distinct from previously characterized roles in peroxisome biogenesis.

LD storage and metabolism is a dynamic process. LD formation occurs even as existing TG stores are broken down by lipases at existing LDs (Hashemi and Goodman, 2015). DGs produced by TG hydrolysis can be re-esterified by DGATs to form new TG that is incorporated into new (initial) LDs (Wilfling et al., 2013). In +Oleate S2 cells, previous RNAi knockdown of *Pex14* suppressed changes in LD volume or number (Figure 3G-H). However, when +Oleate cultured cells were then transferred to Lipolytic conditions, *Pex14* RNAi knockdown reduced LD volume and increased LD number, characteristic of fragmentation that occurs when lipases are active (Figure 3G-H). In addition, larger LDs were observed in cells in Lipolytic conditions compared to +Oleate conditions. Thus, it is possible that the LDs observed in cells treated with *Pex14* RNAi represent new LDs formed from re-esterification of produced by TG lipolysis. In addition, larger LDs were observed in cells in Lipolytic conditions compared to +Oleate conditions. Thus, this increase in LD volume in Lipolytic conditions is likely the result of re-esterification of DG to TG at existing LDs. In other organisms, DGAT isoforms are localized to LDs (McFie et al., 2011) and the FATP1-DGAT2 complex facilitate LD expansions (Xu et al., 2012). Further supporting a role for Pex14 promoting TG storage, LDs in cells overexpressing both *Pex14* and *Hsl* were significantly larger than those found in cells overexpressing *Hsl* alone (Figure 5G-H). This would lead to elevated levels of DG as endogenous Bmm hydrolyzed TG. Given that DG can be re-esterified to TG at the LD surface by an LD-localized isoform of DGAT2 (Stone et al., 2009) in other species, if a similar mechanism is conserved in *Drosophila*, this would cause LD TG stores to be maintained, increasing increase in LD volume. *Diacylglycerol O-acyltransferase 2* (*Dgat2*) is an apparent *DGAT2* homologue, although no functional studies have been performed to confirm similar function, so this possibility is not currently testable. Functional testing of this model awaits functional and biochemical characterization of Dgat2 in *Drosophila*.

The relative increased in TG (glycerol) levels in Lipolytic cells treated with *Pex14* RNAi (Figure 3I), supports a model that Pex14 at LDs suppresses lipase activity. This model is also supported by observations that LD volume in the fat body and overall TG content is significantly reduced when *Pex14* is knocked down by RNAi in the larval fat body (Figure 1E). The role of Pex3 and Pex13 in this process also need to be further examined. Pex13 is predicted to interact directly with Pex14 (Schell-Steven et al., 2005). *Pex13* RNAi knockdown in the larval fat body also caused reduced fat storage and reduced survival on minimal nutrition food supplemented with excess lipid (Figure 1I). The effect of Pex13/Pex14 on larval survival is most likely related to the effect on lipid storage as RNAi knockdown of *Pex1*, which would strongly affect peroxisome biogenesis had little effect on survival when animals were fed a high-fat diet (Figure 1I). Larvae consuming lard food would have increased levels of circulating fatty acids secreted from cells of the gut and stored in fat body (Musselman and Kuhnlein, 2018). *Pex14 RNAi* treatment likely upsets the regulatory balance at LDs in the fat body leading to lipotoxicity. *Pex16* RNAi knockdown blocked Pex14 association with both LDs and peroxisomes (Figure 4D). Pex16 and Pex19 shown previously to have roles in inserting Pex14 into the membrane of peroxisomes or PPVs (Aranovich et al., 2014; Kim et al., 2006; Sacksteder et al., 2000). Conversely, loss of Pex19 leads to an increased proportion of Pex14 localized to LDs in +Oleate cultured cells (Figure 4B) Previous studies have shown that *Drosophila Pex19* mutant larvae have elevated activity of cytoplasmic (non LD-associated) lipases, mitochondrial dysfunction and lipotoxicity (Bulow et al., 2018), consistent with the role we identified for Pex19 in directing Pex14 to LDs which would contribute to systemic defects in lipid metabolism.

Localization of the Pex3, Pex13, and Pex14 to LDs occurs within 24h after cells are placed in +Oleate conditions. Pex14 in the LD proteome is largely synthesised during the first 24h (Figure 3Q-S). Further, we find that LD-associated Pex14 is distinct from markers of mature peroxisomes (Figure 2E) or from mitochondria (Figure 4C). The C-terminal region of Pex14, Pex14^117-280^ is sufficient to mediate LD association in +Oleate cultured cells (Figure 4L). However, LD association is abrogated if the Pex14 C-terminal region does not include the transmembrane domain (Figure 4M). Pex14 is a known bilayer transmembrane protein (Azevedo and Schliebs, 2006; Will et al., 1999) so it is extremely unlikely that the transmembrane domain (Figure 4G) could be inserted into the LD phospholipid monolayer (Fujimoto and Parton, 2011). Thus, the requirement for the TM domain (Figure 4, G-M), the co-recruitment of Pex3 and Pex13 (Figure 3 V-W), the similar phenotypes associated with knockdown of Pex3, Pex13 and Pex14 in the fat body (Figure 1A) and direct TEM visualization of PEX14 localization (Figure 6K-M) supports a model whereby Pex14 at LDs is associated with PPVs.

Pex3, Pex13, and Pex14 are all bilayer membrane spanning proteins inserted into PPVs, in a process that likely requires Pex19 (Götte et al., 1998; Pinto et al., 2006). When inserted into the peroxisome membrane, the Pex14 C-terminal domain faces the cytoplasm (Barros-Barbosa et al., 2019; Reuter et al., 2021). One possible model is that Pex14 at the LD surface is embedded within vesicles and the C-terminal of Pex14 interacts with an LD-resident protein, or that the C-terminal of Pex14 that can interact directly with the LD surface. However, analysis of the C-terminal region of Pex14 did not identify sequence motifs like Class I or Class II LD interacting proteins (Kory et al., 2016). Immuno-TEM of NRK cells showed that LD-associated PEX14 is associated with small vesicles adjacent to LD population (Figure 6K-L). These vesicles are approximately 40-50 nm in diameter, within the range of the size predicted for PPVs (van der Zand et al., 2012). One class of mammalian PPVs is defined by the presence of Pex3 and Pex14 in the membrane (Agrawal and Subramani, 2016; Schrader and Pellegrini, 2017; Sugiura et al., 2017). Further examination as the path taken within the cell Pex3, Pex13, and Pex14 from synthesis in the cytosol to the LD surface in +Oleate cells is needed to determine at what stage these proteins are diverted from the canonical peroxisome biogenesis pathway to LDs.

Supporting the model that the C-terminal of Pex14 associates with the LD surface via interaction with other LD-associated proteins is the observation that changes in the amount of perilipins affects LD localization of Pex14, as well as Pex3 and Pex13. Overexpression of *Lsd-1* blocked the localization of Pex3, Pex13, and Pex14 to the LD surface in S2 cells (Figure 5I, M-N), while elevated Lsd-2 levels promoted Pex14 association with LDs (Figure 5J). A major mechanism regulating association of proteins with the LD surface is inter-molecular competition for limited space as the LD grows with addition of TG and shrinks with TG lipolysis (Kory et al., 2015). As the LD surface shrinks during TG lipolysis proteins are preferentially removed from the LD surface to create space for proteins like lipases (Kory et al., 2015). The exclusion of Pex14 from LDs when Lsd-1 levels are elevated (Figure 5I) suggests this is occurring. As overexpression of *Lsd-2* does not affect Pex14-LD localization, the effect of *Lsd-1* overexpression on Pex14 is likely specific, rather than a general effect. Spatial analysis of the positioning of proteins at surface of individual LDs is needed to define the relative contributions of protein-protein interaction to recruitment of Pex14 (or Pex3 or Pex13) to LDs. Phosphorylated Lsd-1 helps recruits Hsl to the LD surface during lipolysis in *Drosophila* (Bi et al., 2012). This is notable as we have observed that overexpression of Pex14 blocks recruitment of Hsl to the LD surface (Figure 5C-D).

A role for Pex14 in promoting TG storage in LDs is also supported by differences in interaction between Pex14 and LD-associated lipases. The major circulating neutral lipid in *Drosophila* is DG which is accumulated in the fat body for energy storage (Heier and Kühnlein, 2018). While Pex14 levels at the LD have little effect on the TG lipase Bmm, Pex14 antagonizes Hsl at the LD (Figure 5A-H). This supports a model whereby recruitment of Pex14 to the LD surface perturbs the interaction between Hsl and Lsd-1, blocking the recruitment of Hsl to the LD. This model is consistent with the antagonistic effects of overexpression of *Lsd-1* on Pex14-LD localization (Figure 5I). Further studies are required to determine the mechanism by which Pex14 perturbs Hsl recruitment to LDs.

Finally, pulse-chase labelling of newly synthesised protein showing that newly synthesized Pex14 is directed to LDs (Figure 3R-S) and the effect of Pex19 loss on enhanced trafficking of Pex14 (Figure 4E-F) to LDs suggests that LD-localized Pex14 in cells transferred to +Oleate conditions comes from a pool of protein newly translated in the cytosol rather than the pre-existing pool associated with mature peroxisomes. This model of newly translated Pex14 being directed to newly formed LDs in +Oleate cells is also consistent with the observed upregulation of *Pex14* upon transfer to +Oleate culture conditions (Figure 2B) and the lack of an effect on other Pex genes on fat body lipid levels when knocked down by RNAi (Figure 1A-F).

It remains unclear when and how newly synthesized Pex14 is trafficked to the LD. Given the absence of peroxisome proliferation observed in S2 cells transferred to +Oleate conditions, may be that Pex14 newly synthesised in the cytoplasm is diverted from peroxisome proliferation to the LD serving to coordinate these two critical organelles needed for cellular lipid homeostasis. As Pex13 and Pex14 are both inserted into PPVs post-translationally via the activity of Pex3, Pex16 and Pex19 (Giannopoulou et al., 2016; Jansen and van der Klei, 2019) the co-localization and/or functional requirement for each in trafficking Pex14 to LDs suggest that this stage is also required. However, when Pex19 is absent, Pex14 localization to LDs is increased; however, this could also be a function of ablation of peroxisome proliferation in general unbalancing the balance between trafficking Pex13 and Pex14 to each organelle. Clearly, additional experiments will be required to clearly elucidate the transport pathway of Pex3, Pex13, and Pex14 to the LD surface.

## Materials and methods

### Cell culture

‘Standard’ conditions for S2 and Pex19KO S2R+ cell culture: Schneider’s Medium (Sigma Aldrich S0146) containing 10% FBS (Thermo Fisher, 12483-012) at 25°C. The Standard culture conditions for Huh7 or NRK cells were Dulbecco’s Modified Eagle’s Medium (Sigma Aldrich D5796) containing 10% FBS at 37°C and 5% CO_2_. Both were supplemented with 100U penicillin per ml and 100μg streptomycin per ml (Thermo Fisher 15140-122). S2, S2R+, Huh7 and NRK cells were passaged in a log phase before they reached confluency. Cultures were not used beyond passage 25. The +Oleate culture conditions used in this study are the same as used previously to study LDs in S2 cells (Guo et al., 2008; Krahmer et al., 2011), These are the same as Standard conditions except that the medium was supplemented with 1 mM oleate (+Oleate, Sigma Aldrich O1008) bound to fatty acid free BSA (Sigma Aldrich A8806). Cells were maintained in 1 mM oleate-supplemented (+Oleate) conditions for 24 or 48h. To induce lipolysis of LD-stored TGs, (‘Lipolytic’ conditions), cells were first cultured for 24h in +Oleate conditions. Cells were then washed 1x in fresh non-supplemented medium and subsequently incubated in medium without FBS or oleate for 24. The +Oleate and Lipolytic culture conditions were shown previously to induce LD biogenesis and subsequent lipolysis were first described in Guo et al. (Guo et al., 2008).

### Generation of a polyclonal antiserum recognizing Drosophila Pex14

A pENTR-D clone of the full-length Pex14 open reading frame (Baron et al., 2016) was transferred to pDEST-17 (Thermo Fisher) using LR ClonaseII (Thermo Fisher 11791-020). This was transfected into BL21-AI *E. coli* (Thermo Fisher C6070-03). Expression of 6xHis-Pex14 was induced in 500 ml cells grown to OD_600_-0.4 at 37°C by addition of 0.2% L-arabinose and further culture at 25°C for 3h. The bacterial cells were lysed by incubation in 8M Urea, and the lysate cleared by centrifugation 20000xG for 30 min at 25°C. The cleared lysate was applied to a 1ml HisTrap column (Cytavia 17524701), using the Akta-Start His-tagged purification protocol (Cytavia). Purified protein was eluted using a stepwise Imidazole gradient. Fractions containing purified Pex14 were combined, placed in dialysis tubing (8000 MWCO, Spectrum 132660), and desalted by buffer exchange in 5L 1x PBS overnight. The protein sample was concentrated in an Amicon Ultra 15 centrifugal filter (Millipore Sigma, UFC900308) to a concentration 1mg/ml and injected into Guinea Pigs (Pocono Rabbit Farms and Laboratories). Partially purified serum was tested for antigen specificity by western blot against purified protein, S2 cells, S2 cells expressing Pex14-GFP fusions and *Pex14* RNAi treated S2 cells.

### Drosophila strains

The *w^1118^* strain was obtained from the Bloomington *Drosophila* Stock Center (BDSC). RNAi lines used include All crosses were performed at 25°C.UAS-dsRNA lines were obtained from Vienna Drosophila Stock Centre: *Pex1*, GD12029v27741, VDRC:v27741;*Pex2*, KK101378, VDRC:v108578;*Pex3*, GD2464v11017, VDRC:v11017; GD2464v12426, VDRC:v109619;*Pex5*, GD14972v42332, VDRC:105654;*Pex12*, GD11036v34671, VDRC:34671;*Pex11a/b*, KK101579, VDRC:v105654;*Pex13*, GD1977v39544, VDRC:v39544;KK100165, VDRC:v108829;*Pex14*, GD2759v42590, VDRC:v42590;*Pex16*, KK107609, VDRC:v110614;*Pex19*, GD11608v22064, VDRC:v22064;, KK108370, VDRC:v100746 or BDSC: *Pex11a/b*, TRiP.HMS02576, BDSC:42883; *Pex12*, TRiP.HMC03536/TM3, BDSC:53308; *Pex13*, TRiP.HMC03099, BDSC:50697; *Pex14*,TRiP. HMC06491, BDSC:79826; *Pex16*, TRiP.HMC04810, BDSC:57495; *Pex19*, TRiP.HMC03104/TM3(Ubi-GFP), BDSC:50702. Driver lines used were: y^1^ w*; P{w^+mC^=r4-GAL4}3 BDSC:33832or Kyoto Drosophila Stock Center: y^1^ w*; P{TubP-GAL4}LL7/TM3(Ubi-GFP), Sb^1^ KDSC 108069. UAS-Stinger GFP:3:r4-GAL4 was used as a control for RNAi knockdown screen crosses. For flies over TM3(Ubi-GFP), non-GFP flies were selected. Fly stocks were maintained on the BDSC standard cornmeal food recipe, unless specified. Fly strains were passaged once per week to prevent overcrowding.

### Cloning

S2 cell expression clones for Myc or FLAG tagged Pex proteins were described previously (Baron et al., 2016). For expression of epitope-tagged lipid-droplet associated proteins, cDNA libraries were made from mRNA harvested from *Drosophila* embryos at 2-4, 4-6, and 10-14h after egg laying and reverse transcribed using oligo-dT primers with a One-Step RT kit (Bio-Rad 1725140). The coding sequence for each gene encoding a gene of interest was amplified from a cDNA template using Phusion High Fidelity DNA polymerase (Thermo Fisher F-530XL). Full-length coding sequences were amplified for N-terminal tagging of proteins, respectively. Pex14 truncations were similarly generated by PCR of bases 349-840, (aa117-280) and 148-280 (aa 442-840). Pex14 1-444 (aa 1-148) was cloned without a stop codon for C-terminal tagging. Blunt-end purified PCR products with a CACC motif at the 5’ end were directionally cloned into the pENTR/D Gateway entry vector by TOPO cloning (Thermo Fisher K240020). The sequences of the inserted regions were verified by Sanger sequencing at the U of Alberta Molecular Biology Facility. These were recombined into pAFW or pAMW destination vectors for N-terminal tagging or pAWM for C-terminal tagging, which are part of the *Drosophila* Gateway Vector Collection, (originally developed by Terence Murphy, Cornell University) using LR ClonaseII.

### Transfections

Plasmids containing a tagged gene of interest were transfected into *Drosophila* S2 cells using Effectene transfection reagent following the manufacturer supplied protocol (Qiagen 301425). S2 cells were passaged 24h before transfection. Approximately 5.0 x 10^5^ cells were transfected with 150 ng of plasmid DNA. Transfected S2 cells were incubated at 25°C for 48-72h before fixation for imaging.

### ^35^S metabolic pulse-chase labelling

To label newly synthesised protein in S2 cells, a formulation of Schneider’s medium (Schneider, 1972) that did not contain L-methionine, L-Cysteine or yeast extract was made from stock chemicals (Sigma-Aldrich). It was supplemented with 10% dialyzed FBS (Thermo Fisher A33820) and 100μl Easy Tag Express ^35^S Methionine/Cysteine (Perkin Elmer NEG 772002MC) at either 0 or 24h after transformation with a 6xMyc-Pex14 (pAMW-Pex14) as described above. 1mM oleate was added at 24h after transformation as described above. For cells where 35S was added at 0h, they were washed in complete Schneider’s medium (Schneider, 1972) at 24h. For cells where ^35^S Methionine/Cysteine mix was added at 24 h, these were washed in Schneider’s medium at 72h. Cells were pelleted at 72h and rinsed with PBS containing Complete protease inhibitor cocktail (Millipore 04693159001) and fractionated as described below. The relative proportion of 6xcMycPex14 was analyzed by immunoprecipitation from each fraction and detected by autoradiography or western blotting.

### Imaging

S2 cells were observed using a Zeiss 63x oil immersion objective (NA = 1.4) on a Zeiss Axio Observer M1 microscope with an ERS spinning disk confocal and a C9100 EMCCD camera (Hamamatsu) using Volocity imaging software (PerkinElmer) or a Zeiss LSM700 confocal and Zen software (Zeiss). Image stacks were captured at 130μm vertical (z) spacing (ERS) or 25nm (LSM700). S2 cells were fixed in 4% paraformaldehyde dissolved PBS and blocked with 3% bovine serum albumin (BSA). Cells were incubated for one hour with anti-FLAG M2 monoclonal mouse primary antibody (Sigma-Aldrich F3165), anti-Myc rabbit primary antibody (Sigma Aldrich SAB4301136), anti-Pex14 Guinea pig primary antibody (Simmonds lab), anti-Abcd3 rabbit primary antibody (Simmonds lab), anti-cytochrome (BD Pharmingen, Clone 7H8.2C12) mouse primary antibody and anti-SKL rabbit primary antibody (Baron et al., 2016). The first two primary antibodies were used at a 1:200 dilution, and the last four were used at 1:1000, 1:500, 1:500 and 1:250, respectively. In cases where 6xMyc-tagged proteins were analyzed relative to peroxisomes, anti-Myc mouse primary monoclonal antibodies 9E10 (obtained from Dr. Paul Lapointe, University of Alberta) or 9B11 (Cell Signalling 2276S) were used at a 1:250 dilution. Primary antibody incubation was followed by incubation with Alexa Fluor 568 anti-mouse goat secondary antibody and Alexa Fluor 488 anti-rabbit secondary antibody, both at 1:2000 dilution. Cells were imaged as above.

To detect LDs, cells were stained with HCS LipidTOX Deep Red at 1:500 dilution for one hour after secondary antibody incubation. For larval fat body staining, fat bodies from third instar larvae were dissected and fixed in 4% paraformaldehyde for 15 minutes. The tissue was rinsed three times in PBS. The tissue was then stained with Nile Red (Thermo Fisher) and 4,6′ - diamidino-2-phenylindole (DAPI) at a 1:1000 and 1:500 dilutions, respectively. Tissues were mounted on slides with Prolong Gold (Thermo Fisher, P36930) mounting medium and imaged. NRK and Huh7 cells were fixed in 4% paraformaldehyde and blocked in 3% BSA for one hour. Cells were incubated in rabbit anti-PEX14 primary antibody (Thermo Fisher PA5-78103) and mouse anti-peroxisome membrane protein 70 / ATP-binding cassette, subfamily D, member 3 (ABCD3) primary antibody (Richard Rachubinski, University of Alberta) at 1:200 dilutions, for one hour. Primary antibody incubation was followed by incubation with Alexa Fluor 568 anti-mouse goat secondary antibody, or Alexa Fluor 488 anti-rabbit secondary antibody (Abcam ab175473 and ab150077), Alexa Flour Cy3 anti-Guinea Pig secondary antibody (Jackson ImmunoResearch 706-165-148), and Alexa Flour 488 anti-mouse secondary antibody, and Alexa Flour 594 anti-mouse secondary antibody, all at 1:2000 dilutions. Nile Red (Sigma Aldrich 3013) or LipidTOX Deep Red (Thermo Fisher H34477) were used at 1:500 dilution to stain LDs. Cells were imaged with a 63x objective lens, as above. The images shown best represent the quantitative data that accompanied them. In Figures 1 -2, images were quantified from six biological replicates. In Figures 3, 4G-J, 5-10 images are representative of three biological replicates. Figure 4 images are representative of four biological replicates.

### Image processing and quantification

Image stacks of individual confocal images comprising the entire cell volume were processed to remove noise and reassign blur using a classical maximum likelihood estimation confocal algorithm provided by Huygens Professional Software (Scientific Volume Imaging) and an experimentally determined point spread function constructed from multiple images of 0.1μm Tetraspeck beads (Thermo Fisher T7279). Three-dimensional-based colocalization analysis using Pearson’s coefficient was performed with Huygens Professional Software (Scientific Volume Imaging. In this case, colocalization is defined as the co-occurrence of two fluorophores. This was quantified using Pearson’s coefficient whereby a value of +1.00 (100%) denotes complete colocalization and 0 (0%) denotes the absence of any colocalization (Adler and Parmryd, 2010). Peroxisome or LD volume and average number of peroxisomes or LDs per cell were calculated using IMARIS v8 (Oxford Instruments). To estimate what percentage of the co-localization signal was due to background fluorescence, measurements were also calculated on images where one channel was shifted 90°, relative to the other (Dunn et al., 2011). In all cases, background co-localization measured in the shifted images were never greater than 10%. For quantification of PEX14 immunolocalization on TEM images, the distance between PEX14-positive signals and LDs was measured and reported based on the proportion of the total PEX14 signal within a given image (Figure 6M). These distances were grouped into six ranges: 0-40 nm, 41-80 nm, 81-120 nm, 121-160 nm, 161-200 nm, and 200+ nm.

Organelle volume and number were measured using the Surfaces function as follows: the specific channel for the appropriate organelle marker was selected. A Gaussian filter was applied by selecting “Smooth” and the surfaces detail was set to 0.1μm. Thresholding was set to “Background Subtraction (Local Contrast)”. The diameter of the smallest organelle signal to be included in the measurement was measured in the “Slice” mode, and the value was inputted into the Surfaces creator below “Background Subtraction”. A surface was created, and the background signal was removed by adjusting the slider in the Surfaces creator. Finally, surfaces were created by selecting the green arrow to perform the appropriate calculations. Values were found under the “Statistics” tab, which gives the total number of surfaces (organelle number). Organelle volume was given by selecting “Average values”.

In Figure 2F, peroxisome number values represent averages based on 10 cells measured from three biological replicates, for a total of 30 cells measured. In Figures 2 E, G-H, peroxisome and LD volume and number values represent averages based on five cells measured from six biological replicates, for a total of 30 cells measured. In Figure 5F, LG volume values represent averages based on six images measured from three biological replicates, for a total of 18 imaged measured. In Figure 5K-L, colocalization values represent averages based on five cells measured from four biological replicates, for a total of 20 cells measured. In Figure 6E-F, colocalization values represent averages based on five cells measured from three biological replicates, for a total of 15 cells measured. For statistical analysis of all colocalization and organelle volume/number data, an unpaired Student’s *t*-test was performed using Prism 7 software (GraphPad).

For Figure 4G-H, spectral imaging and linear unmixing were performed using Zen software (Zeiss). The two dyes, Alexa Flour 594 and Cy3 were imaged using Lambda Mode with wavelengths between 521.0 to 630.0nm and the number of channels were adjusted until width was 10nm. In the Unmixing tab, Auto find/ACE (Automatic Component Extraction) was selected to extract 7 spectral components in the acquired image. The appropriate ACE channel were selected and deconvolved as above. The DAPI, neonGreenand LipidTOX deep red images were acquired separately using standard filter settings.

### dsRNA treatments

dsRNA amplicons were made from an existing template library (Foley and O’Farrell, 2004). RNA was amplified using a T7 RNA Polymerase (Thermo Fisher, EP0111). S2 cells were passaged 24h prior to dsRNA treatments. Cells were treated with dsRNA using Effectene Transfection Reagent (Qiagen 301425) to enhance uptake. S2 cells were incubated for 72h at 25°C before further processing. A scrambled dsRNA amplicon was used as a control (Forward primer sequence: GTGAAGAGGTCAGAGGCCTG; Reverse primer sequence: ACAGTCTAGCGTTCCTTGAGG.

### qRTPCR analysis

RNA was isolated from S2 cells using the RNeasy Plus Mini Kit (Qiagen 74134). RNA was reverse transcribed using the Maxima H minus system (Thermo Fisher K1681).

Quantification of each transcript was performed using Perfecta SYBR Green FastMix (QuantaBio 95118) and an Eppendorf MasterCycler RealPlex2. All samples were measured in triplicate and calculations were made relative to Ribosomal Protein L30 (RpL30) expression. Primers used for each of the target genes were previously experimentally validated pairs reported in FlyPrimerBank (Hu et al., 2013). For all qRTPCR experiments, values reported are averages based on three biological replicates. Statistical significance was determined by unpaired Student’s *t-*test or one-way ANOVA test using Prism 7 software (GraphPad).

### RNA-SEQ and analysis

Total RNA was isolated from S2 cells cultured in Schneider’s or Schneiders +Oleate culture conditions using an RNeasy Plus Mini Kit. RNA integrity was verified using an Agilent RNA Nano assay (Agilent Genomics 5067-1511). Ribosomal RNA was subtracted from samples using a Ribo-Zero Gold rRNA Removal Kit (Illumina 20040526). Libraries were prepared using a NEBNext Ultra RNA Library Prep Kit and NEB Next Multiplex Oligos (New England Biolabs E7530L and E7335L). Library quality and size distribution was confirmed by running an Agilent High Sensitivity DNA assay (Agilent Genomics 5067-4626), and the average size of library inserts was verified to be 290 - 300 base pairs. 10pM of each of the libraries were loaded onto an Illumina MiSeq v2 300 cycle kit (2 x 150 cycles, paired-end reads, MS-102-2003). Each culture condition was analyzed in triplicate. Paired-end reads were aligned to the *Drosophila melanogaster* genome (6.28 release) HiSat2 (Kim et al., 2015). Individual read counts were mapped to specific genes using HTSeq (Anders et al., 2015). Reads with less than one count per million in at least three samples were filtered out (Pertea et al., 2016). Differential analysis was performed using a pipeline that correlated EdgeR (Robinson et al., 2010) and DESeq2 (Love et al., 2014) modified from the SARTools pipeline (Varet et al., 2016). Transcripts that were found to have differential expression in both alignment and differential expression models (padj>0.1) were considered for subsequent analysis.

### Larval buoyancy assays

Approximately four days after egg-laying, late 3^rd^ instar larvae were removed from their vials, rinsed in sterile PBS and suspended in a 12% sucrose solution, as per Reis *et al*. (Reis et al., 2010). Larvae were scored by their propensity to float in the sucrose solution. In each trial, 10 larvae were analyzed. Three biological replicates were performed for each sample, for a total of 100 larvae analyzed. For analysis of Pex14 GD2759v42590, 10 individual replicates were performed, and statistical significance was measured by unpaired Student’s *t*-test using Prism 7 software (GraphPad).

### TG / glycerol quantification

At approximately 4 days after egg-laying, 3^rd^ instar larvae were removed from their vials and rinsed in sterile PBS. Lysates were made by homogenizing tissue in 5% NP-40 in distilled, deionized water. Samples were heated at 80°C for 5 minutes, cooled to room temperature, and centrifuged to remove any insoluble material. TG measurements from each sample were made using a Triglyceride Assay Kit (Abcam, ab65336), as per the manufacturer’s instructions. Fluorometric detection was made at 587 nm using a BioTek Synergy 4 plate-reader with Gen 5 software. TG measurements were made relative to the protein concentration of each sample, measured using the Pierce BCA Protein Assay Kit (Thermo Fisher 23225). Lysates were made from 10 larvae for each trial. The values reported are averages from three biological replicates. Statistical significance was measured by unpaired Student’s *t-*test using Prism 7 software (GraphPad).

The glycerol content of cell culture media was quantified using a Glycerol Assay Kit (Sigma Aldrich MAK117) according to the manufacturer’s protocol. S2 cells were pelleted by centrifugation, and the resulting Schneider’s medium was removed. Each sample was diluted 1:1000 in water. The assay was performed per the manufacturer’s instructions, and end-point fluorescence was measured at 587 nm in a BioTek Synergy 4 plate-reader with Gen 5 software. Glycerol measurements were made relative to the protein content in each sample, measured using a Pierce BCA Protein Assay Kit (Thermo Fisher 23225). For protein measurements, cells were lysed in Mild Lysis Buffer (20 mM HEPES pH 7.0, 50 mM NaCl, 1 mM EDTA, 0.5 mM EGTA, 10 mM DTT, 1.0% Triton X-100, protease inhibitors), and protein measurements were taken, as per manufacturer’s instructions. Colorimetric absorption was measured at 562 nm using a BioTek Synergy 4 plate-reader with Gen 5 software. The values reported are based on averages from six biological replicates. Statistical significance was determined using any unpaired Student’s *t*-test (Prism 7 software, GraphPad).

### Larval survival assay

For the lipotoxicity experiments shown in Figure 1 I, early 3^rd^ instar larvae were transferred to standard cornmeal food to holidic food (Piper et al., 2014) or lard food prepared as holidic food with the addition of lard at 22.2 g/L. For each trial, 10 3^rd^ instar larvae from each genetic cross were transferred to holidic food or lard food. The values shown are averages from five individual genetic crosses, for a total of 50 larvae examined from each genetic cross.

### Subcellular fractionation

LDs were isolated from transfected S2 cells, as described (Krahmer et al., 2011; Krahmer et al., 2013). In brief, cells from a T25 flask were pelleted, washed in cold PBS, and resuspended in 2ml of buffer (200 mM Tris/HCl pH7.5, 2mM magnesium acetate) with protease inhibitors (Roche). The cells were lysed using a cell homogenizer and a 10-μm ball bearing (isobiotec). The lysates were then cleared by centrifugation at 1,000x g for 10 minutes. 1 ml of cleared lysate was adjusted to 1.08 M sucrose, and a step gradient of sucrose was layered on top with 2ml of 0.27M sucrose, followed by 2ml of 0.135M sucrose. Finally, one ml of 0M sucrose buffer was layered at the top. Samples were spun at 100,000x g for 90min at 4°C in an ultracentrifuge using an SW41 rotor. The floating LD fraction was isolated, and the proteins within the fraction were precipitated by methanol: chloroform extraction, (Wessel and Flügge, 1984). In brief, 2 ml of methanol and 500μL of chloroform was added to 500μL of LD fraction isolate. The mixture was vortexed and centrifuged at 9,000x g for 10 seconds. 1.5 ml of ddH_2_O was added, and the mixture was again vortexed and centrifuged at 14,000x g for one minute. The top aqueous layer was removed, and an additional 2 ml of methanol was added and vortexed. The sample was centrifuged at 20,000x g for 5min to pellet the protein. The methanol was carefully removed, and the protein pellet was dried. The dried protein pellet was resuspended in 30μL of gel sample buffer, boiled, and size separated by SDS-PAGE.

### Immunoblotting

Protein samples were boiled in gel sample buffer for 5 minutes, size separated by SDS-PAGE, and transferred to a nitrocellulose membrane (Bio-Rad 1620112). Membranes were blocked in Odyssey Blocking Buffer (LI-COR). For subcellular fractionation experiments, membranes were incubated with rabbit anti-Lsd-2 primary antibody (obtained from Dr. Michael Welte, University of Rochester) and mouse 9B11 (Cell Signalling 2276S) anti-MYC antibody. For the metabolic labelling experiments blots were re-probed with Rabbit anti SKL (Richard Rachubinski). For NRK cell lysates, membranes were probed with rabbit anti-PEX14 primary antibody (Thermo Fisher) and E7 mouse anti-β-tubulin primary antibody developed by Klymkowsky was obtained from the Developmental Studies Hybridoma Bank, created by the NICHD of the NIH and maintained at The University of Iowa, Department of Biology, Iowa City, IA 52242. Membranes were then probed with Alexa Fluor anti-rabbit A680 secondary antibody and Alexa Fluor anti-mouse A790 secondary antibody (Abcam ab175773 and ab175783). Membranes were visualized using an Odyssey Infrared Imaging System (LI-COR), and the band were quantified using Odyssey software (LI-COR). Western blots were representative of three independent biological replicates.

## Acknowledgements

### Funding

This work was supported by a Discovery Grant from the Natural Sciences and Engineering and Research Council of Canada to AJS.

## Supplemental material

**Supplementary Figure 1.**
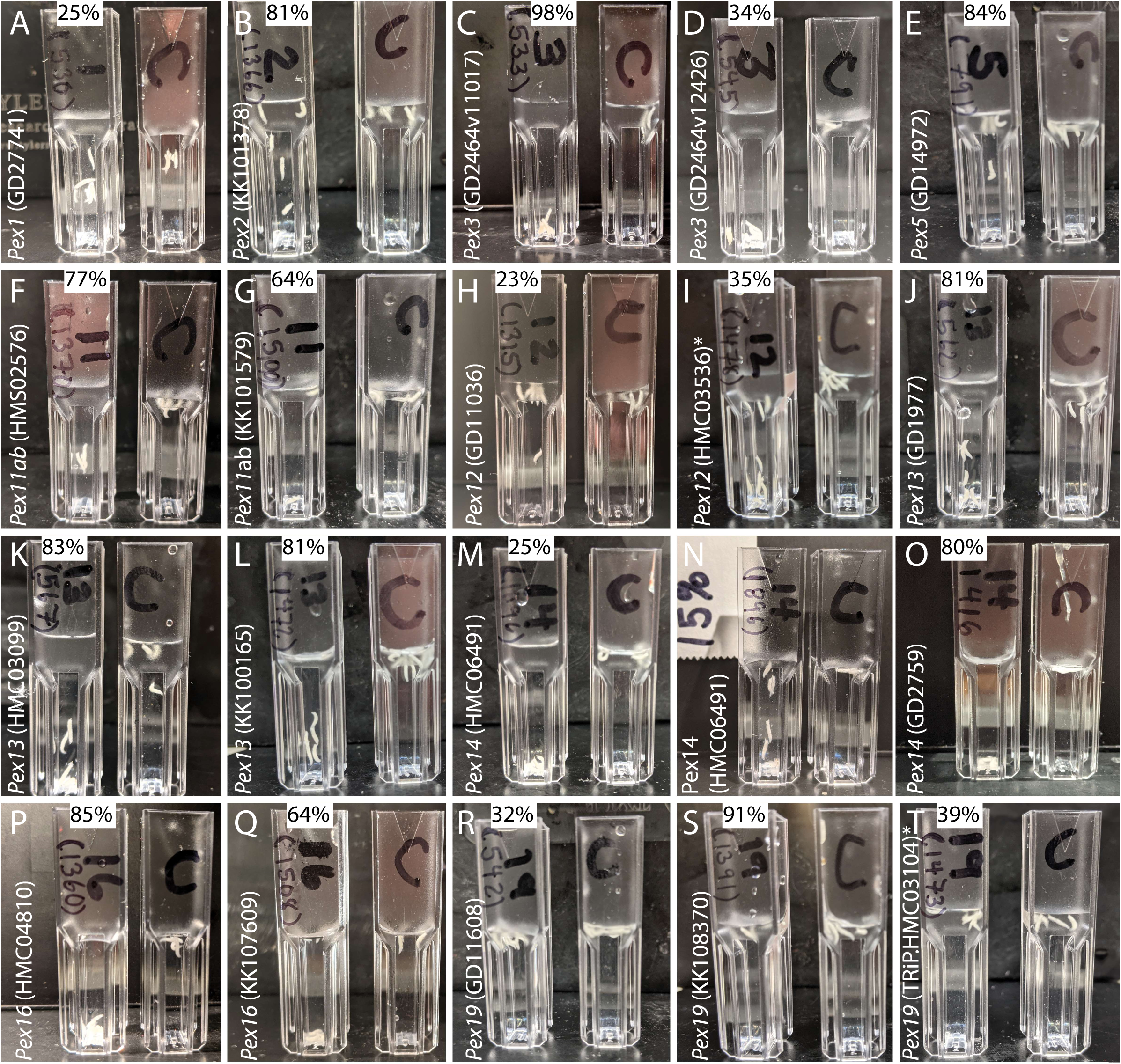
A screen for the effect of Pex knockdown on the fat body. Representative images from a small-scale screen for the effect of *Pex* gene knockdown on lipid storage in larvae with. The r4-GAL4 driver line and UAS-transgenes expressing double stranded RNA targeting each Pex gene were used (See Figure 1 B-C). The value at the top of each image indicates the efficiency of RNAi knockdown confirmed by qRTPCR in larvae where the UAS-RNAi transgene was expressed ubiquitously via Tub-GAL4. Larvae with normal fat storage float in 12% sucrose. For each image, the cuvette shown on the left is the knockdown experiment and, on the right, (labelled C) control r4-GAL4 larvae suspended in a 12% sucrose solution, except for (N) where a 15% sucrose solution was used. The unique ID for each RNAi transgene is provided. The number in brackets under the Pex gene number on each cuvette indicates a serial number used for experimental blinding. The position of the larvae in each cuvette was recorded in terms of quartiles representing distance from the surface of the sucrose solution to the bottom.

Supplementary Table 1-Complete RNA SEQ data

Supplementary Table 2-GO classification of RNAs differentially expressed in S2 cells cultured in Standard versus +Oleate conditions.

